# Deep learning model for the prediction and classification of protein toxins across all domains of life

**DOI:** 10.1101/2021.06.29.450401

**Authors:** Javier Caceres-Delpiano, Roberto Ibañez, Simon Correa, Michael P. Dunne, Pedro Retamal, Leonardo Álvarez

## Abstract

Toxins are widely produced by different organisms to disrupt the physiology of other organisms, and support their own existence. Their study is useful to understand protein evolution, environmental adaptation and survival competition. *In-silico* predictions of toxic proteins can support empirical frameworks, and help in the safety measurements needed for various industrial related processes. Some *in-silico* methods are slow, hard to implement or lack taxa representation in their training datasets. Here we present a deep learning model to classify protein toxins, through the use of Convolutional Neural Networks (ConvTOX). ConvTOX is able to accurately identify toxic proteins across the domains of life, with accuracies over 80% for animal and plant toxins, and over 50% for bacterial toxins. Moreover, ConvTOX is able to generalize the identification of differences among toxin types, such as neurotoxins and myotoxins, and to accurately identify structural similarities between different protein toxins. ConvTOX overcomes limitations from previous models by being able to predict toxin proteins from across all domains of life, and by not being limited to only short toxin peptides. Limitations are still clear in terms of lower accuracies for specific phylogenetic groups (such as bacterial toxins), but still this works presents itself as a one step forward for the universal use, classification and study of toxic proteins.

## Introduction

Toxins are biological substances that can cause negative effects on other organisms, both externally and internally, with important pathological effects that can impact on physiological functions such as immune response, irritation of mucosa, organ failure, and even death^1^. It is not exactly known how many toxins exist, but as bioassay technologies improve, the availability of toxicity data increases and new hypotheses can be developed and tested. Unfortunately, only a limited number of toxins have undergone comprehensive toxicological evaluations due to the limitations of traditional toxicity testing^2^. Traditional analysis methods fail to effectively process large and complex data, such as toxin classification datasets, presenting both a challenge and an opportunity to explore new ways of data processing.

Recently, *in silico* methods have been successfully developed to predict protein toxicity, where identification of potential toxic proteins is generally performed by scanning protein sequence patterns, or performing multiple sequence alignments, against different databases, such as animal toxins^3,4^, spider venoms^5^ and microbial pathogens^6^, using BLAST, and Hidden Markov State models^7^, locating conserved toxic motifs (tox-bits), and serving as the basis of general toxin classifications that allow the prediction of whether a certain substance could be harmful. However, toxin evolution poses a limitation to these kinds of methods, since protein toxins usually evolve from proteins with a physiological function, and thus specificity of these methods is limited. Additionally, these approaches do not possess the required sensitivity to identify non-sequence related structural similarities^8,9^.

Deep learning (DL), a machine learning method that takes advantage of deep neural networks (DNN), has become a widely used approach to establish relationships and patterns from high-dimensional data obtainable from proteins such as sequence, three dimensional atomic coordinates, and secondary structure^10,11,12^. In the area of protein toxins, some machine learning methods have been developed based on the use of support vector machines (SVM), such as ToxClassifier^13^, that can classify venom and non-venom proteins, encoding sequences as combinations of two amino acids, Hidden Markov Models (HMM) of tox-bit motifs, and BLASTp searches against positive and negative venom databases^13^. Another example is TOXIFY, which uses recurrent neural networks (RNN) based on matrices encoding Atchley factors per amino acid to classify animal toxins^14^. These methods have shown to be useful, but only cover toxins of a narrow spectrum of phylogenetic groups, leaving out toxins from plant and prokaryotic origin. In order to make accurate predictions, deep learning algorithms must be able to generalize from the training data to all unseen observations. This approach requires data with reliable sampling of the observations that we are interested in. Therefore, the larger the taxonomic representation, the easier it will be for the model to generalize to unseen data. Thus, it is worth including information of toxins from across all domains of life in the development of new predictive methods, possibly allowing the identification of important relationships that might exist among toxic proteins between different phylogenetic groups.

In this article we present a new neural network model with supervised learning that has been trained with information on toxic proteins from organisms across all domains of life, which can distinguish toxic from non-toxic proteins. We evaluate our method by means of clustering analyses of known toxic protein sequences, and benchmarking against similar machine learning models. To complement these evaluations, we also tested the model’s capability to clusterize and organize different protein toxins into toxin types and structure-based groups.

## Methods

### Training datasets

The training dataset for the prediction models were composed as follow:

- For our positive dataset we included all protein sequences obtained from UniProtKB, classified by Keyword [KW-0800] as toxin, and with a length from 4 to 500 residues. Redundant sequences were filtered at 90% sequence identity. All reviewed and non-reviewed sequences were included. This resulted in a total of 27,869 protein sequences classified as toxic.
- For our negative dataset, random sequences were selected from UniProtKB, using the search term ‘NON Keyword [KW-0800]’, and with a length from 4 to 500 residues. Redundant sequences were again filtered at 90% sequence identity. A total of 40,532 sequences were selected.

A random 90% subset from the sequence data was used as positive and negative training sequences, while a remaining 10% was used as a test set for model validation.

### Protein sequence encoding

A crucial step for model training and testing is the encoding of amino-acid sequences through an encoding scheme, in order to assign a numerical representation for each amino-acid. The encoder takes a protein sequence P of length L as X = (x1, x2, …, xL) and encodes it into a sequence of vector representations of the same length Z = (z1, z2 …, zL). To encode all protein sequences as a high-dimensional numerical representation, we used Tasks Assessing Protein Embeddings (TAPE), which was pretrained on a dataset of over 31 million protein sequences, focused on different prediction tasks^11^.

### Sequence classification model

For the protein classification model, we used encoded sequences as input (see below). Four hidden layers of Convolutional 1D Neural Networks over the sequence length dimension were used, with 768 neurons each and kernel size of 5. The output of the Convolutional layers was a tensor of dimension 768. Finally we added two linear layers to reduce dimensionality from 768 to 512 and then from 512 to 2, which allows predicting toxicity as binary classes. A learning rate of 0.01 was used, and cross-entropy was used as loss. The training was run for 100 epochs, and the accuracy and loss were recorded for every epoch. The final model was selected as the model with best accuracy (epoch 41 in our case), and it was used to estimate the probability that a given protein sequence was classified as a toxin. In some cases, a softmax function was applied to our binary output to estimate the probability of proteins belonging to different classes such as: non-toxin, highly non-toxin, highly toxin, and toxin.

### Validation and benchmarking

To benchmark our model, we used 249 proteins with ≤500 amino acids, not included in the training and validation set of ConvTOX. This set was composed of 85 animal toxins, 40 plant toxins and 124 bacterial toxins, and was used to compare the performance against state-of-the-art models. ToxClassifier and TOXIFY were the selected models to compare performance, since these two methods included proteins of similar sizes as ConvTOX. Other models, such as ToxinPred, only included short toxin peptides no larger than 35 residues.

For validation and testing, we collected two different protein sets, not contained in the training set. The first one included different toxin types collected from UniprotKB using different keywords: 75 neurotoxin proteins (with Keyword KW-0528), 85 enterotoxin proteins (with Keyword KW-0260), 62 myotoxins proteins (with Keyword KW-0959), 46 dermonecrotic toxin proteins (with Keyword KW-1061), 18 cell-adhesion impairing toxic proteins (with Keyword KW-1217) and 19 G-protein coupled receptor impairing toxic proteins (with Keyword KW-1213). The second set included 141 toxic proteins with known structure extracted from the Protein Data Bank (PDB).

## Results

### Model validation

The overall accuracy of the model (from now on called ConvTOX) was calculated at every epoch over the test and validation datasets. After training, the best epoch, in terms of those accuracies, was selected. At epoch 41, the model showed a 94.63% accuracy on the test set and 94.80% accuracy on the validation dataset, which was the one selected as the best.

To visualise how ConvTOX learnt, a vector representation of each sample was generated by the model (Fig. 1A), and then a dimensionality reduction was performed to further reduce the dimensions into two classes: toxic and non-toxic proteins. Figure 1B shows the sample distributions in the test set by means of a two-dimensional t-distributed stochastic neighbor embedding (t-SNE)^15^. Positive and negative protein samples are separately clustered, indicating that the model is effective for the classification of protein toxins. Percentages of 95.6% and 97.5% of sensitivity and specificity, respectively, were reported by the model.

**Fig. 1.**
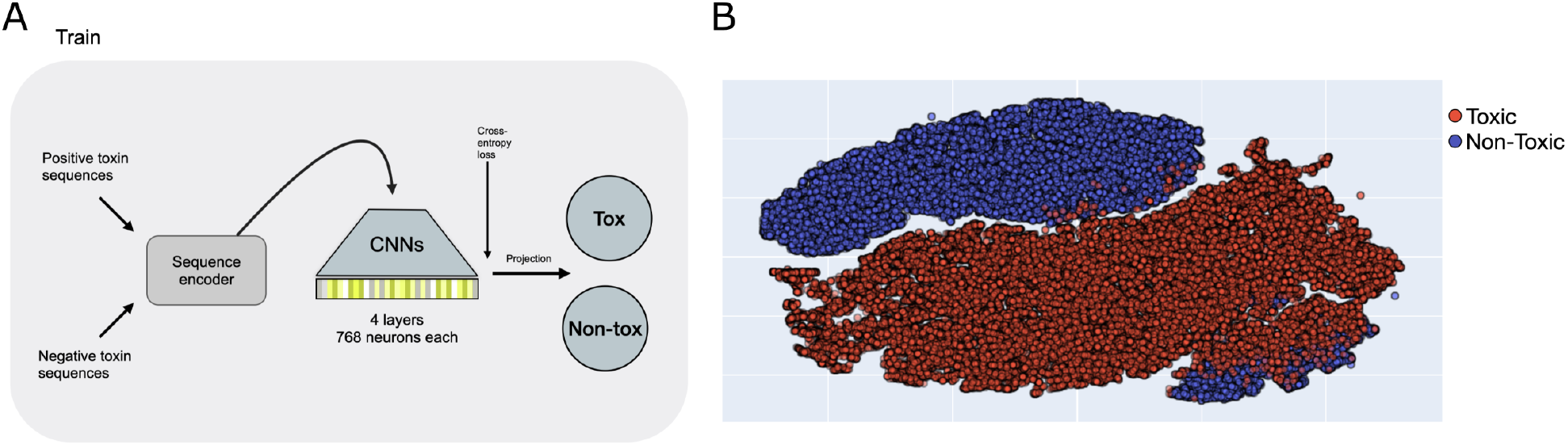
Model structure for toxins classification. **(A)** Workflow of information in the training of ConvTOX, including input data, information encoding, and training with the use of convolutional neural networks (CNNs) with cross-entropy as loss. The model was used to estimate the probability of a protein being a toxin. **(B)** t-SNE visualization of protein sequences in the test set colored by ground-truth classification of toxic (red) and non-toxic (blue) proteins.

### Benchmarking against state-of-the-art models

Since our model was trained with toxic protein sequences from different kingdoms (i.e. animal, plant and bacterial toxins), we decided to follow a similar benchmarking approach to previous studies^13,14^. For this, 249 proteins not included in the training and test set, composed of 85 animal toxins, 40 plant toxins and 124 bacterial toxins, were used. Even though this approach to comparison with other methods is not totally fair, since TOXIFY and ToxClassifier were trained with animal toxins only^13,14^, we can not rule out that both methods can still contain information that let them generalise to other kingdoms. In the case of animal toxins, ToxClassifier outperforms TOXIFY and ConvTOX, with a true-positive percentage of 95.29%, corresponding to 81 out of 85 correctly classified animal toxins (Fig. 2). Concerning the other two models, we found TOXIFY to perform comparably to ConvTOX, with true-positive percentages of 85.88% (73 out of 85 correctly classified) and 83.52% (71 out of 85 correctly classified), respectively (Fig. 2). For the case of other taxa, ConvTOX outperforms ToxClassifier and TOXIFY. In the case of plant and bacterial protein toxins, ConvTOX presented a true-positive percentage of 87.5% (35 out of 40) and 56.45% (70 out of 124), respectively (Fig. 2). ToxClassifier is unable to classify plant and bacterial toxins, while TOXIFY only achieves percentages of 5.0% and 3.2%, respectively. It is worth noting that the low performance in the prediction of bacterial toxins might be due to the lower representation of these types of toxins in the training dataset, which only represents less than 10% of the total set.

**Fig. 2.**
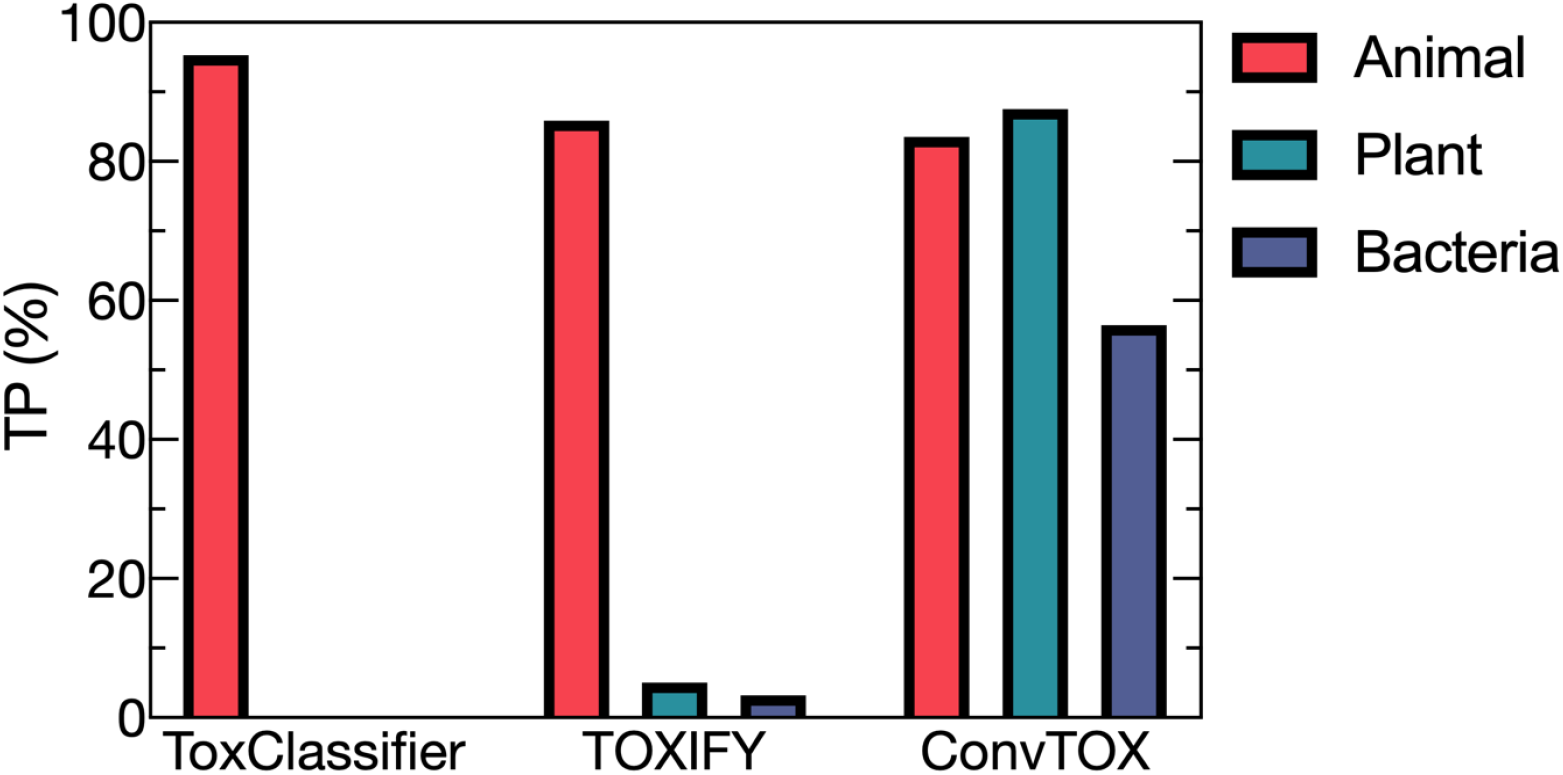
Benchmark for toxic proteins against different classification models. Benchmark against ToxClassifier and TOXIFY, based on true-positive percentages. Correctly classified toxin percentages were calculated for toxins from different kingdoms, such as animal (red), plant (green) and bacterial (blue) toxins.

### Generalization of toxin information by ConvTOX

A key challenge in machine learning methods is the generalization of models to regions outside their training domain. Deep learning models in particular often struggle in this regard^16^. Even though ConvTOX was trained in a supervised manner on a wide sample of toxic and non-toxic proteins from different kingdoms, it is expected that there will exist toxins unseen by the model that may differ significantly from those in the training and test sets. It is therefore important to check that the model is able to capture general toxin features, such as toxin types and structural similarities.

To test this, a set of different toxin types was collected, including reviewed neurotoxins, enterotoxins, myotoxins, dermonecrotic toxins, cell-adhesion impairing toxins and G-Protein coupled impairing toxins, from UniProtKB (see *Validation and benchmarking* section in methods). We found that the model was able to capture crucial information in order to generate clear protein clusters. A t-SNE projection for a set of different toxins showed meaningful toxin type clusters (Fig. 3A). Even though we trained the model to classify toxic from non-toxic proteins, the model is able to extract crucial features from several toxic protein sequence types that let us discriminate similarities and differences between them, and those results correlate with what is found in the literature.

**Fig. 3.**
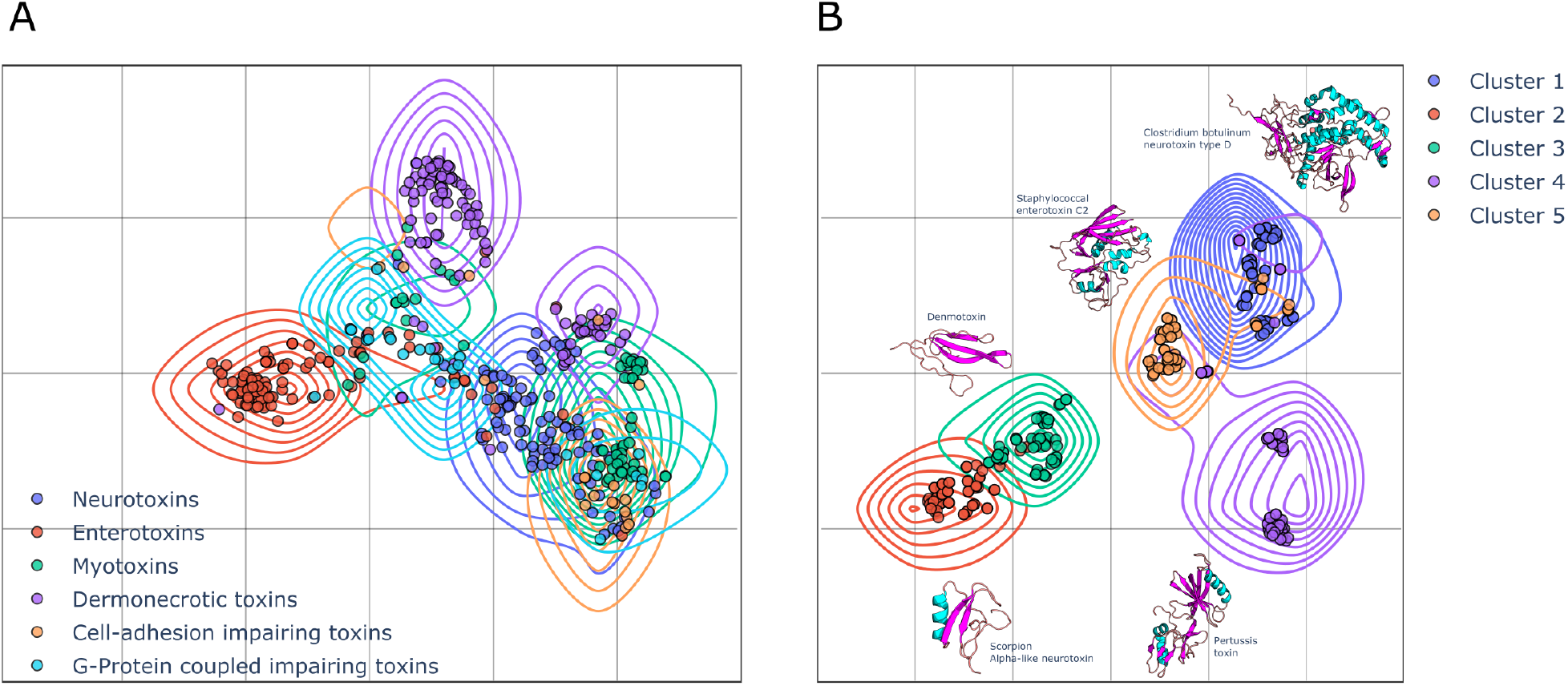
Toxin types and structural classifications. **(A)** t-SNE visualization of different types of toxic protein sequences colored by ground-truth toxin types, such as neurotoxins, enterotoxins, myotoxins, dermonecrotic, cell-adhesion impairing, and G-protein coupled impairing toxins. **(B)** t-SNE visualization of toxins with known structure extracted from PDB, with representative 3D protein structures for each well-defined cluster.

Finally, to assess how structurally related proteins are represented by ConvTOX, we examined its ability to cluster structurally similar toxins that do not necessarily present high sequence identity or come from the same organism, but which might share similar three-dimensional structures. For this, a set of protein toxins with known three-dimensional structure was collected from PDB (see *Validation and benchmarking* section in methods). These protein sequences were fed to ConvTOX to test its capability to cluster similar structures, and to possibly find interesting toxin relations. As shown in figure 3B, different clusters are clearly formed, and a structural representative per cluster is shown. For luster 1, most structures were related to neurotoxins such as Botulinum and Tetanus neurotoxins. Cluster 2 was fully associated to scorpion toxins, while cluster 3 was related to venomous elapid snakes toxins such as neurotoxic proteins of the three-finger toxin superfamily (e.g. Denmotoxin, Irditoxin and Erabutoxin). Cluster 4 was related to AB5 toxins such as Pertussis toxin from *Bordetella pertussis*. Lastly, cluster 5 showed proteins such as Staphylococcal enterotoxin and Streptococcal pyrogenic exotoxin.

## Discussion

In this work we show the capabilities and potential applications of a new toxin classification model called ConvTOX. This model uses deep learning to take advantage of known protein toxins, encoding their sequences into high-dimensional vectors that contain the crucial information needed for classification using convolutional neural networks. ConvTOX also presents itself as an important alternative to slower BLAST-based methods and SVM models^13^.

Compared to state-of-the-art models, ConvTOX makes clear progress in a key limitation for those models, such as TOXIFY^14^ where only high accuracy is obtained in the classification of venom proteins across eumetazoans. Here, we overcome those limitations by including information for different taxa, improving prediction accuracy for those types of protein toxins. Moreover, the model is able to generalize to unseen observations, such as to clearly identify protein toxins of different classes, and to group structurally similar protein toxins, approaches that have not been previously studied for other models. Clear limitations are still seen in the estimation of bacterial toxins, mostly due to the fact that toxins for other taxa outnumber the bacterial toxins used in this work. As mentioned in other studies^13,14^, classification and prediction of protein toxins can crucially be improved by increasing the amount of empirical verifications for different protein sequences. We believe that these models will play a key role in the support of these empirical approximations, speeding up the verification process for proteins of unknown toxicity. Finally, this type of model can be used to support existing frameworks in the safety assessment process for proteins under various industrial applications, for example as a high-throughput approach to accelerate the filtering of multiple proteins before cytotoxicity assays, or to even support the evaluation of GRAS (Generally Recognised As Safe) status.

